# Seeing speech: The cerebral substrate of tickertape synesthesia

**DOI:** 10.1101/2022.09.19.508477

**Authors:** Fabien Hauw, Mohamed El Soudany, Charlotte Rosso, Jean Daunizeau, Laurent Cohen

## Abstract

Reading acquisition is enabled by deep changes in the brain’s visual system and language areas, and in the links subtending their collaboration. Disruption of those plastic processes commonly results in developmental dyslexia. However, atypical development of reading mechanisms may occasionally result in ticker-tape synesthesia (TTS), a condition described by Francis Galton in 1883 wherein individuals “see mentally in print every word that is uttered (…) as from a long imaginary strip of paper”. While reading is the bottom-up translation of letters into speech, TTS may be viewed as its opposite, the top-down translation of speech into internally visualized letters. In a series of functional MRI experiments, we studied MK, a man with TTS. We showed that a set of left-hemispheric areas were more active in MK than in controls during the perception of normal than reversed speech, including frontoparietal areas involved in speech processing, and the Visual Word Form Area, an occipitotemporal region subtending orthography. Those areas were identical to those involved in reading, supporting the construal of TTS as upended reading. Using dynamic causal modeling, we further showed that, parallel to reading, TTS induced by spoken words and pseudowords relied on top-down flow of information along distinct lexical and phonological routes, involving the middle temporal and supramarginal gyri, respectively. Future studies of TTS should shed new light on the neurodevelopmental mechanisms of reading acquisition, their variability and their disorders.

**Significance statement:** Some individuals, whenever they are hearing speech, see vividly in their mind’s eye the corresponding words in written form, as mental subtitles. This unusual condition, termed ticker-tape synesthesia (TTS), far from being purely anecdotal, actually touches on the core of the brain mechanisms of normal and impaired reading acquisition. Through 3 fMRI experiments, plus brain connectivity analyses, we propose an in-depth study of a single individual with such ticker-tape synesthesia. We propose that TTS, a situation in some sense symmetrical to developmental dyslexia, reflects an inverted flow of information through the reading system, such that speech is automatically translated into internally visualized letters. Future studies of TTS should shed new light on the neurodevelopmental mechanisms of reading acquisition.

## Introduction

Reading acquisition is enabled by deep functional and anatomical changes in the brain’s visual system and language areas, and in the links that subtend their collaboration (Dehaene et al., 2015; Dehaene-Lambertz et al., 2018; López-Barroso et al., 2020; Thiebaut de Schotten et al., 2014). Whenever those delicate plasticity processes are compromised, various sorts of developmental dyslexia may result (D’Mello and Gabrieli, 2018; Friedmann and Coltheart, 2016). However, atypical development of reading mechanisms may occasionally result in another condition, nearly opposite to dyslexia, one which has not received sufficient scientific attention.

The polymath Francis Galton noted in 1883 that “some few persons see mentally in print every word that is uttered and they read them off usually as from a long imaginary strip of paper, such as is unwound from telegraphic instruments” (Galton, 1883). This unusual phenomenon, dubbed by Galton ticker-tape synesthesia (TTS), once purely anecdotal, actually touches on the core of the brain mechanisms of normal and impaired reading. Indeed, while reading is the translation of visual letters into speech, TTS may be viewed as its exact opposite, i.e. the translation of speech into internally visualized letters, respectively the “inducer” and “concurrent” in synesthesia parlance (Holm et al., 2015; Ward and Simner, 2020).

During word reading, letters and their order are first identified in the Visual Word Form Area (VWFA), a reproducible sector of the left ventral occipitotemporal (VOT) cortex, whose lesion yields pure alexia (Cohen et al., 2000; Dehaene et al., 2015; Dejerine, 1892). Orthographic information is then broadcast to language areas, giving access to the associated sounds (the “phonological route”) and meaning (the “lexico-semantic route”) (Coltheart, 2012; Perry et al., 2007; Taylor et al., 2013). Reciprocally to this “bottom-up” spread of information, brain imaging revealed that in literate individuals, language areas exert “top-down” influences upon orthographic representations. Thus, attended speech activates the VOT cortex including the VWFA (Cohen et al., 2021; Ludersdorfer et al., 2016; Yoncheva et al., 2010), and language areas send different top-down influences to the VWFA during the reading of real words and of pseudowords (Sharoh et al., 2019). In this framework, as first hypothesized by Holm et al. (2015), TTS would reflect exceptionally powerful and automatic top-down influences from speech processing areas onto the VWFA. Using functional MRI (fMRI) in a man in whom TTS was triggered by both real words and pseudowords, we assessed the hypothesis that TTS would essentially upend the reading system to generate letters from sound.

We implemented this hypothesis as the following four specific predictions. (1) TTS should activate perisylvian language areas involved in speech processing and phonology-to-orthography translation, and VOT regions subtending orthography. (2) Regions involved in TTS should overlap with those involved in reading. (3) Assuming that TTS reflects intense topdown flow of information in the reading system, dynamic models should reveal an increased drive of the reading system by speech input. (4) Due to the overarching distinction between the lexical and phonological routes, top-down influences should follow different paths within the reading system depending on whether synesthesia is triggered by words or pseudowords.

## Materials and Methods

### Participants

All information concerning the TTS participant, MK, is reported in the Results section. Two groups of healthy controls participated in the study. One group (G1) included 22 participants, aged 20-36 years (median 28 years, 10 men). The other group (G2) included 14 older controls, aged 51-75 years (median 64 years, 5 men). All participants were native French speakers, right-handed according to the Edinburgh Inventory (Oldfield, 1971), had no history of neurological or psychiatric disorders. The research was approved by the institutional review board of the INSERM (protocol C13-41), and all participants provided informed written consent. All methods were performed in accordance with the Declaration of Helsinki.

### Experiment 1: Normal and reversed speech perception

#### Stimuli and procedure

The Red Riding Hood story was recorded with a female voice, and a time-reversed version of the recording was derived by playing it backward. MK and group G1 received a pseudo-random alternation of 6 blocks of normal and 6 blocks of reversed speech (block duration 15 seconds), separated with silent rest periods, for a total of 5 min and 12 s. Group G2 received a slightly shorter version (4 normal and 5 reversed blocks, for a total of 4 min and 15 s). Participants were simply asked to pay attention to the story.

#### MRI acquisition

For MK and G1, multi-echo (ME) fMRI images (Posse, 2012) were acquired on a Siemens 3T MAGNETOM Prisma scanner, with a 20-channel receive-only head coil: TR 1.3 s; multi-TE 15, 34.24, 53.48 ms; flip angle 68°; voxel size 3 × 3 × 3 mm; 48 slices. An anatomical T1-weigthed image was acquired: TR 2.3 s; TE 2.76 ms; flip angle 9°; voxel size 1 × 1 × 1 mm). For G2, single-echo sequences were acquired on a Siemens 3T MAGNETOM Verio scanner, with a 20-channel receive-only head coil: TR 3 s; TE 25 ms; flip angle 90°; voxel size 2 × 2 × 2.5 mm; 49 slices. An anatomical T1-weigthed image was acquired: TR 2.3 s; TE 4.18 ms; flip angle 9°; voxel size 1 × 1 × 1 mm).

#### Preprocessing

For MK and G1, using afni_proc.py, all echo timeseries were slice-time corrected, motion correction was computed on the first echo and applied to all echos, and echo images were merged using t2smap.py from the meica.py toolbox provided with AFNI: http://afni.nimh.nih.gov/afni/. Then, using SPM12 (Wellcome Institute of Imaging Neuroscience, London, UK) implemented in Matlab, functional time-series were co-registered to the anatomical volume, normalized, resampled to 3 mm cubic voxels, and spatially smoothed (6 mm FWHM). For G2, time-series were preprocessed only with SPM12, following the same steps: slice-timing, realignment, co-registration to the anatomy, normalization, resampling to 3 mm cubic voxels, smoothing (6mm FWHM).

#### Regions of interest

In addition to the whole-brain approach, we restricted some analyses within two a priori defined regions of interest (ROI). First, we used a 8 mm radius sphere centered at the peak of the VWFA (MNI −44 −50 −14), as identified in Dehaene et al. (2010). Second, we created a ROI covering left-hemispheric language areas and VOT cortex (Supplementary Figure S1), by merging the opercular and triangular parts of the inferior frontal gyrus, the supplementary motor area, the precentral, fusiform, inferior parietal, supramarginal, angular, superior and middle temporal, and inferior temporal gyri, as defined in the AAL 3 atlas (Rolls et al., 2020).

#### Statistical analysis

One subject was excluded due to technical issues during acquisition. For single-subject analyses, we used a general linear model (GLM) including regressors for normal and reversed speech, convolved with the canonical SPM hemodynamic response function (HRF), plus 6 motion parameters, and high-pass filtering (128 s cutoff). For second-level analyses, we used two sample t-tests to assess differences between MK and control participants, with age as a covariate. We set both the first- and second-level voxel-wise cluster-forming threshold to p<0.001, and the cluster-wise threshold to p<0.05 corrected for multiple comparisons across the whole brain or within ROIs. In the text and in Tables, we provide Z-scores for contrasts of interest at peak activation voxels.

### Experiment 2: Visual localizer

#### Stimuli and procedure

Only MK participated in this experiment. He was presented with an alternation of blocks of pictures (8s per block) and periods of rest (7.8 s per period). Each stimulation block included eight pictures from one of the following categories: printed words, numbers, faces, houses, tools, body parts. Each picture was displayed for 600 ms and followed by a 400 ms blank screen. During rest and inter-trials intervals, a black central fixation cross was presented to minimize eye-movements. The experiment included 10 s of initial rest, followed by 30 blocks of pictures (six for each category) and 30 periods of rest. Blocks were presented in pseudorandom order to maximize the variety of transitions between conditions while avoiding repetition of the same condition in successive blocks. Participants were asked to press a button with their right thumb whenever a picture was identical to the previous one, which was the case for 20% of stimuli (1-3 repetitions/block).

#### MRI acquisition, preprocessing, and statistical analysis

Functional images were acquired on a Siemens 3T MAGNETOM Verio scanner equipped with a 64-channel receive-only head coil: TR 1022 ms; TE 25 ms; flip angle 62°; voxel size 2.5 × 2.5 × 3 mm; 45 slices. Preprocessing was the same as for single-echo data in Experiment 1. Statistical analysis and thresholding followed the same method as in Experiment 1, apart from the definition of regressors of interest: here we defined one regressor for each of the 5 categories of stimuli, plus a regressor for responses to targets.

### Experiment 3: Listening to words and pseudowords

#### Stimuli and procedure

MK and the G1 group of control subjects participated in this experiment. We generated a list of 64 words and 32 pseudowords matched in number of phonemes and syllables. Real words included 32 high- and 32 low-frequency words, whose activation pattern did not differ, and which were averaged in all analyzes. Stimuli were synthetized with a female French voice, and equated for maximum amplitude. Stimuli had a mean duration of 644 ms, with no difference between words and pseudowords.

Stimuli were organized in 48 mini-blocks of 8 stimuli each. Mini-blocks were grouped in 8 consecutive blocks, each composed of 2 mini-blocks of PW and 4 mini-blocks of words. The run started and ended with a 15 seconds silence, and an additional 15 seconds pause was present between blocks. The total duration was about 12 minutes. Subjects had to detect the occasional target pseudoword “tatatata”, pronounced by the same voice as stimuli.

#### MRI acquisition, preprocessing, and statistical analysis

Acquisition and preprocessing were the same as for multi-echo data in Experiment 1. Univariate analyses and thresholding followed the same method as in Experiment 1, apart from the definition of regressors of interest: here we defined regressors for low-frequency words, high-frequency words, pseudowords, targets and responses to targets. As controls were all younger than MK, an age regressor would have absorbed any difference between the two. Hence no age regressor was included in second-level analyses. Moreover, multivariate pattern analyses (MVPA) were conducted using The Decoding Toolbox (Hebart et al., 2015), in order to decode words from pseudowords. To train and test the support vector machine, we performed 8 leave-one-block-out cross validation folds, across the 8 blocks of the experiment. Decoding was performed across the volume activated by the averaged words and pseudowords (voxelwise p<0.01), using a 12mm radius searchlight. Decoding performance around each voxel was quantified as the “area under the curve” (AUC) of signal-detection theory. In order to assess the significance of decoding accuracy in individual participants, we computed a permutation test, with a total of 128 permutations. We report voxels with better-than-chance decoding performance (p<0.05), with moreover an ad-hoc minimum cluster size of 50 voxels.

### Dynamic Causal Modelling (DCM)

We performed deterministic DCM analysis on Experiments 1 and 3, using the DCM toolbox implemented in SPM12. First, a GLM was defined for each experiment, with one regressor modeling all auditory stimulations, plus a parametric modulator taking the value +1 or −1 for normal and reversed speech in Experiment 1, and the value +1 or −1 for words and pseudowords in Experiment 3. Time series were extracted from 4 ROIs identified in Experiments 1 to 3: (1) The mid superior temporal gyrus ROI (STGm) was defined as a 6mm radius sphere centered on the activation peak of reversed speech > rest in controls in Experiment 1; (2) The supramarginal gyrus ROI (SMG) was the cluster, identified in the contrast normal speech > reversed speech in MK > controls in Experiment 1; (3) The middle temporal ROI (MTG) was the cluster of effective lexicality decoding in MK in Experiment 3; (4) The VWFA was defined as a 6mm radius sphere centered on the activation peak of words > faces and houses in MK in Experiment 2. The VWFA and STG ROIs were defined as spheres because activation clusters were large and extended to adjacent irrelevant regions. We specified a set of a priori constraints on the connections between the 4 regions (defining possible model “families”), and on the modulations acting on those connections, as fully presented in the Results section. In MK, within those constraints, we successively identified for each experiment, (1) the most likely set of intrinsic connections between ROIs, and (2) the most likely pattern of modulation of those connections by experimental factors. In brief, we pooled models into model families, each of which shares the same intrinsic connectivity pattern. Within each family, different models then corresponded to different connectivity modulation patterns.

## Results

### A case of tickertape synesthesia

MK was a 69 years old man, right-handed, native French speaker, a still active engineer and researcher in the field of radar technology, with no significant medical history. For as long as he could remember, when listening to speech, he had been simultaneously perceiving the words in their written form. Five years before he first contacted us, he discovered incidentally that this phenomenon was quite unusual. TTS was triggered by speech perception, irrespective of whether it was somebody else or MK himself speaking, of whether he could see the face of the speaker or not, of whether he was himself speaking overtly or covertly “in his head”.

During TTS, MK felt that he perceived distinctly each and every letter. Mental images of words did not occupy a well-defined position in external space, but were rather “in his head”; still they looked black, and could be in upper or lower case. Size was “normal, as when you’re reading”. He felt that he perceived about one word at a time. The illusion appeared immediately upon speech onset and persisted briefly after speech ended. TTS was unaffected by the voice, the gender, or the emotion of the speaker, nor by the content of the utterance. It occurred even with unattended speech, such as from neighbor travelers on a train. MK could not voluntarily inhibit TTS, and found parasitic speech quite disturbing when he was reading, although this may be a common nuisance. MK enjoyed listening to wordless music, which generated no TTS. He had no other associated type of synesthesia. His daughter and a niece were subject to the same phenomenon (Tilot et al., 2018).

### Experiment 1: The correlates of tickertape synesthesia during speech perception

We first scanned MK with fMRI while he was listening to continuous speech (the Red Riding Hood story), to time-reversed speech as a control for low-level auditory perception, and while he was at rest, and his activations were compared to neurotypical controls (see Methods). As expected, MK reported that listening to normal speech induced vivid synesthesia, while reversed speech did not generate orthographic imagery.

In controls, reversed speech minus rest activated the bilateral mid-superior temporal gyrus (STG), inferior frontal gyrus (IFG) and supplemental motor area (SMA), and the left precentral gyrus (Figure 1 and Table S1). The contrast of normal minus reversed speech additionally activated the bilateral left-predominant IFG and superior temporal sulcus (STS) from the temporal pole to the angular gyrus (AG), the left precentral cortex, the bilateral SMA and precuneus, plus the VOT cortex along the lateral occipitotemporal sulcus anterior to MNI Y=-62 (Figure 1 and Table S1).

**Figure 1.**
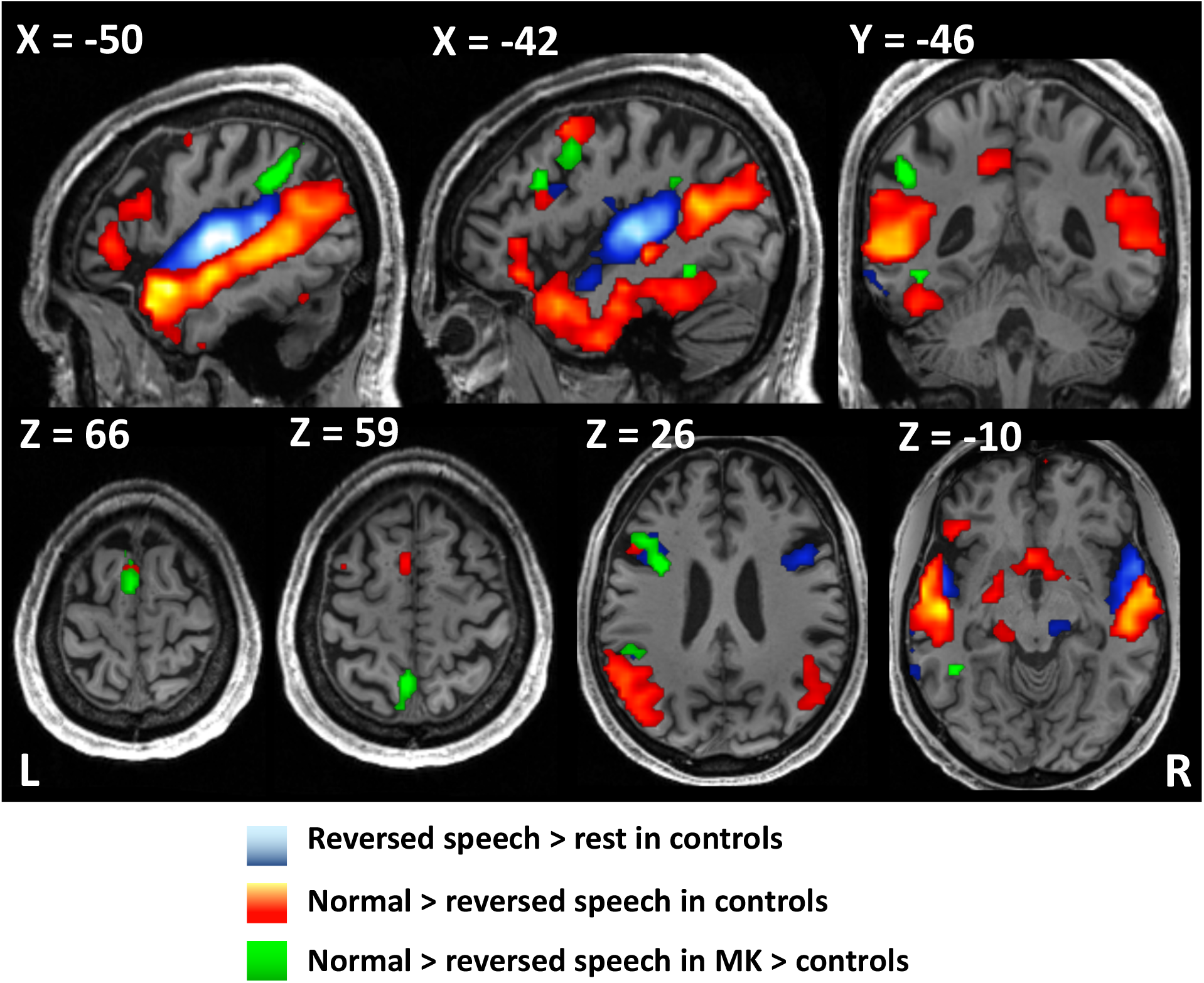
Experiment 1: Correlates of tickertape synesthesia (TTS) during speech perception. Activation in control participants by reversed speech minus rest (blue), and by normal minus reversed speech (hot colors). The latter included left-predominant perisylvian areas usually involved in speech comprehension, plus the left ventral occipitotemporal cortex. When listening to normal speech, MK was subject to TTS and showed a larger difference between normal and reversed speech than controls in the left SMG, IFG, VWFA, precuneus, and SMA (green). TTS, tickertape synesthesia; IFG, inferior frontal gyrus; SMG, supra-marginal gyrus; VWFA, visual word form area; SMA, supplementary motor area.

We then compared MK to controls. In order to identify the functional correlates of TTS, we compared the contrast of normal minus reversed speech in MK relative to controls (Figure 1 and Table 1). MK’s activation was stronger than in controls in a set of left-hemispheric regions: the IFG (MNI −33 5 26, Z=5.59; MNI −42 20 29, Z=5.02), the supramarginal gyrus (SMG; MNI −48 −43 32; Z=6.47), precuneus, the SMA, plus the fusiform gyrus close to the usual location of the VWFA (MNI −42 −46 −10; Z=4.25, corrected within an anatomically defined VOT region of interest; see Methods). We did not find differences between MK and controls in low-level visual cortex. For brevity, we will refer to this set of regions as the TTS network in subsequent results. We defined an anatomical mask including all language areas and the left VOT cortex (see Methods). The TTS network was almost entirely included in those regions (Supplementary Figure S1), in agreement with our first prediction.

**Table 1.**
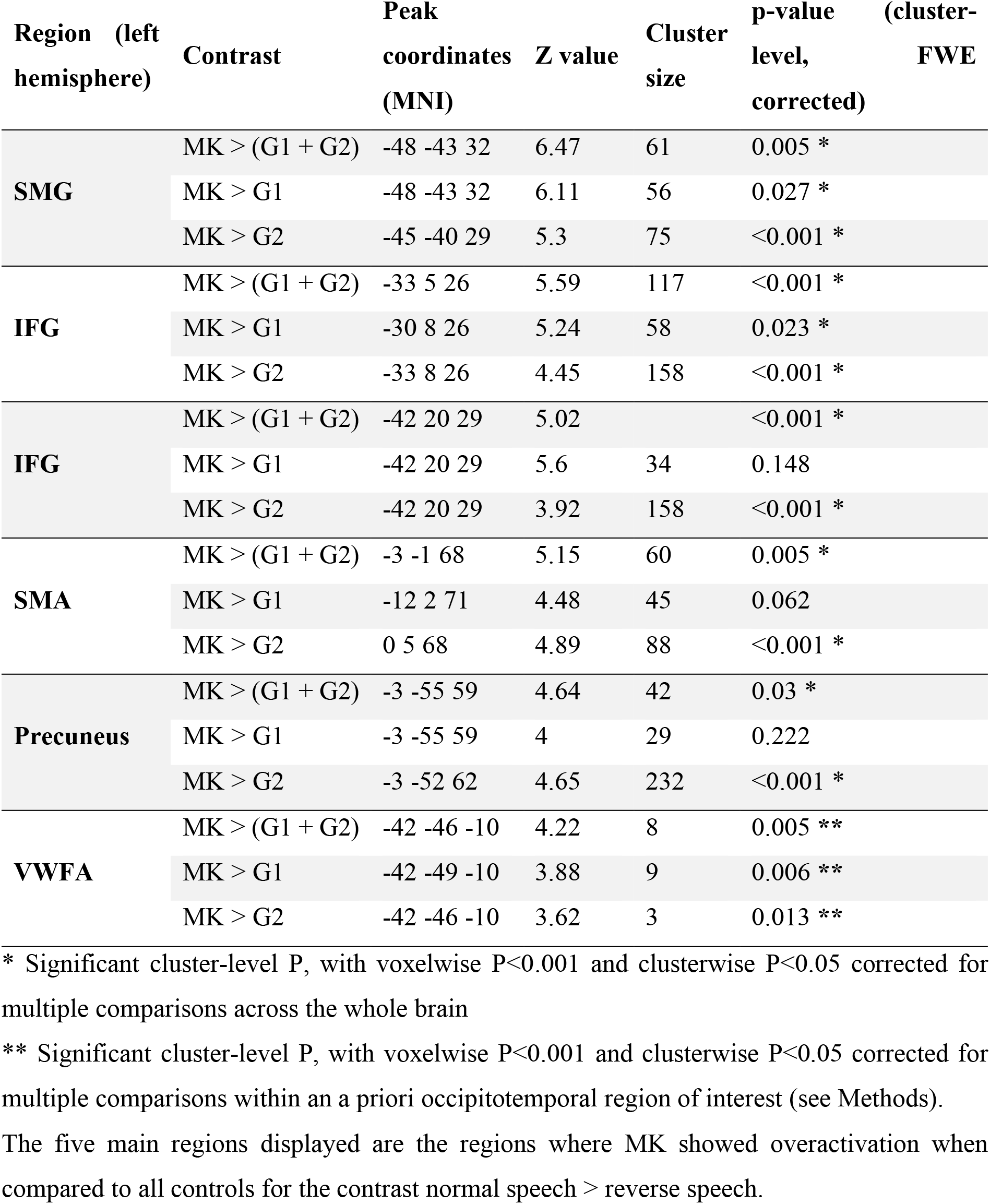
Experiment 1: Regions more activated in MK than in controls when listening to normal > reversed speech.

In order to determine whether MK’s over-activations were within the range of variation of control participants, we compared the contrast of normal minus reversed speech in each control subject minus all others. While 14 controls showed some significant differences from the group (excluding the cerebellum), those over-activations were randomly distributed all over the brain, with little overlap with the language and VOT masks (overlap range 0-58 voxels; mean = 11, as compared to MK’s overlap of 232 voxels; Crawford t-test: t(34)=11.2; p<0.001).

The two control groups G1 and G2 were scanned on different magnets, and G1 controls were younger than MK. In order to make sure that the overall differences which we observed between MK and controls were not due to age and scanner differences, we compared separately MK > G1 and MK > G2, and found that the main differences were remarkably replicated across both analyses, using the same statistical thresholds as in the global comparison (Figure S2). The MK > G1 comparison showed overactivation in the left SMG, IFG and VWFA. The MK > G2 comparison showed overactivation in the left SMG, IFG, SMA, precuneus and the VWFA (see Table 1). Moreover, we directly compared the two control groups. We found stronger activation for G1 > G2 in bilateral superior temporal sulci (STS), and stronger activation for G2 > G1 in the right prefrontal cortex (Figure S3). Those regions had no overlap with the areas distinguishing MK from controls.

Finally, the opposite contrast of normal minus reversed speech in all controls minus MK showed no activations. Activation by reversed speech minus rest and by normal speech minus rest did not differ between MK and controls.

In summary, this experiment supported our first prediction by showing abnormally strong activation during TTS, both in perisylvian frontoparietal language areas (the IFG and SMG), and in the left VOT at the usual location of the VWFA.

### Experiment 2: Tickertape synesthesia as reverse reading

Beyond proposing that TTS should involve both the perisylvian and the VOT regions, we predicted that TTS-related activations should overlap with regions activated during word reading. Particularly, is the VOT component of the TTS network identical to the VWFA, as usually defined by word-specific visual activations? In order to delineate his reading network, MK was scanned while viewing images of words, faces, houses, tools, body parts, numbers, and during simple fixation (see Methods).

We first characterized the mosaic of VOT regions showing category preferences (Figure 2 and Table S2). MK had a fully normal pattern of specialization, as documented in a host of previous studies (see e.g. Deen et al., 2017; Malach et al., 2002; Monzalvo et al., 2012). Subtractions of each of the 3 categories words, faces and houses minus the other 2 showed the left VWFA (MNI −45 −52 −10; Z>8), the right fusiform face area (FFA), and the bilateral parahippocampal place area (PPA), respectively. Subtractions of tools and of body parts minus the previous 3 categories showed bilateral lateral occipital cortex (LOC) activations.

**Figure 2.**
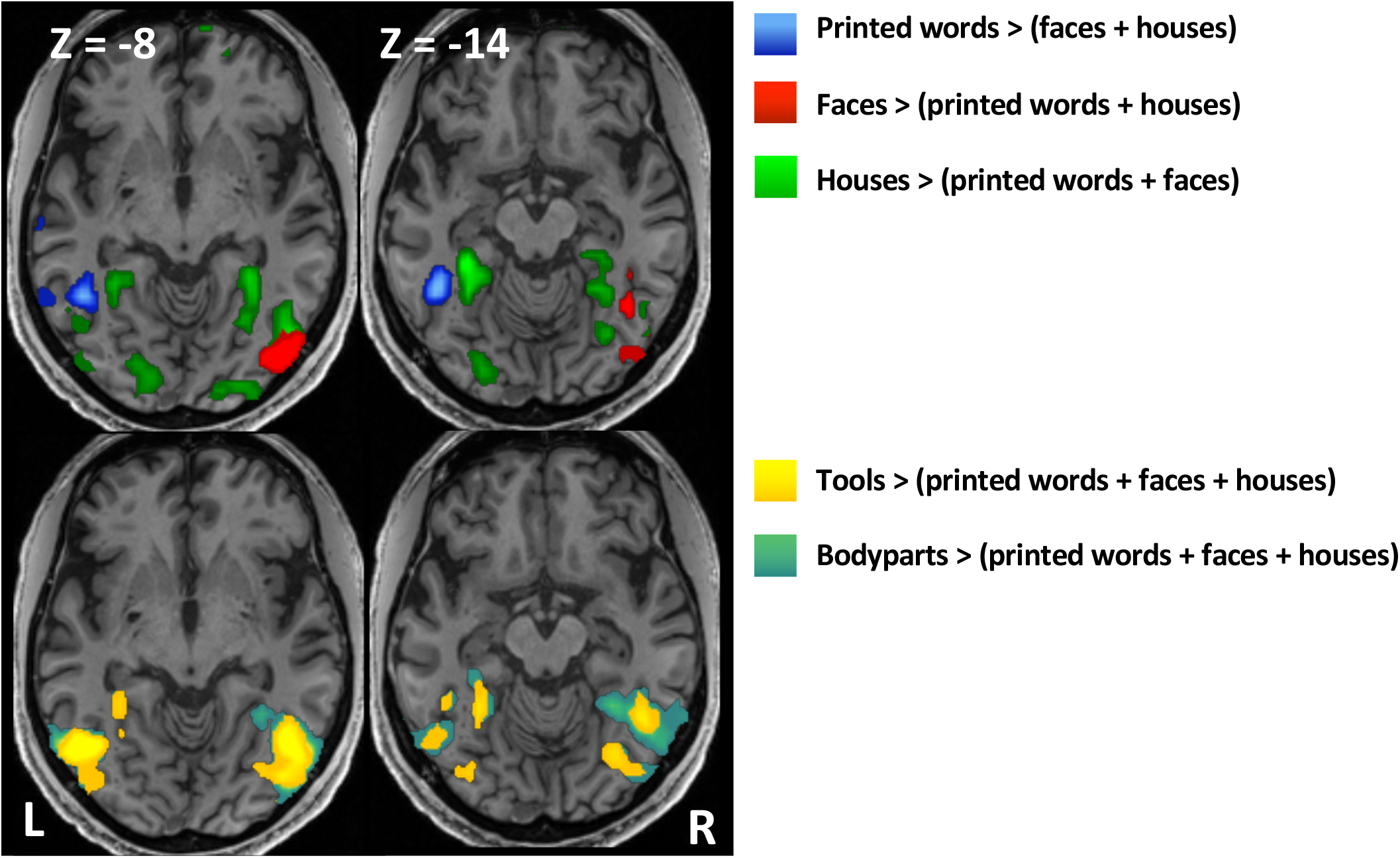
Experiment 2: Category-selective activations by visual objects. MK’s VOT cortex showed a typical mosaic of category preference. Upper row: the VWFA for words (blue), the FFA and OFA for faces (red), the PPA for buildings (green). Bottom row: lateral occipital activations for tools (yellow) and body parts (blue-green). VOT, ventral occipito-temporal; VWFA, visual word form area; FFA, fusiform face area; OFA, occipital face area; PPA, parahippocampal place area.

Crucially, the VOT area which was overactivated during TTS in Experiment 1 was identical to the VWFA defined here on the basis of visual stimulation (Figure 3A, white arrow). Beyond the VWFA, this overlap extended to all components of the TTS network (SMG, IFG, precuneus and SMA), which were also significantly activated here during reading (Figure 3A).

**Figure 3.**
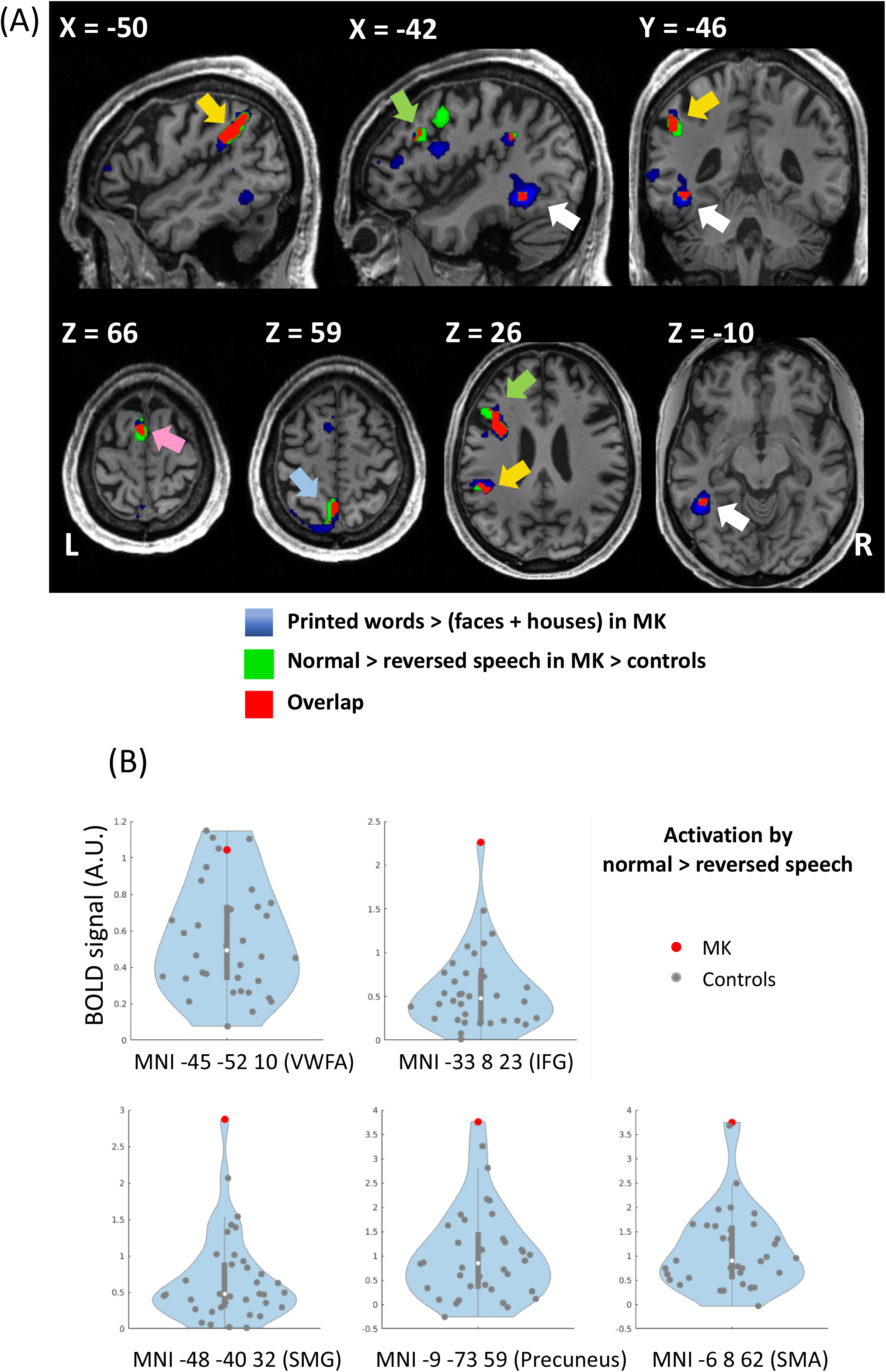
Experiments 1 and 2: Tickertape synesthesia (TTS) as inverted reading. (A) In MK, areas over-activated during TTS in Experiment 1 (green) and areas selectively activated by printed words in Experiment 2 (blue) showed perfect overlap (red), in the VWFA (white arrow), SMG (yellow arrow), IFG (green arrow), SMA (pink arrow) and precuneus (light blue arrow). (B) Plot of individual activation intensity for the contrast of normal > reversed speech (Experiment 1), in MK (red dots) and controls (grey dots), at the peaks of MK’s word-selective activations as identified in Experiment 2. In the IFG, SMG, precuneus and SMA, MK exceeded all of the 35 controls. TTS, tickertape synesthesia; VWFA, visual word form area; IFG, inferior frontal gyrus; SMG, supra-marginal gyrus; SMA, supplementary motor area.

Finally, in order to further compare MK to controls while avoiding statistical double dipping (Kriegeskorte et al., 2009), we defined ROIs of the TTS network on the basis of the current experiment, and plotted data from Experiment 1. In a 10 mm radius sphere centered on peaks of MK’s activation by words minus faces and houses, we computed the average activation by the contrast of normal > reversed speech from Experiment 1, across the 10% most activated voxels (Figure 3B). In the SMG, IFG, precuneus and SMA, MK exceeded all of the 35 controls, with a significantly stronger activation (Crawford’s t-tests: all Ps ≤ 0.001). At the VWFA, 4 controls showed higher activation than MK, who was marginally stronger than the group of controls (p=0.055).

In summary, we observed an extensive overlap of the activations related to TTS and to word reading, supporting our hypothesis that TTS resulted from an atypical operation of the cerebral reading system.

### Experiment 3: Tickertape synesthesia for words and pseudowords

The reading system is thought to associate a phonological route mapping orthography to sounds on the basis of statistical regularities, and a lexico-semantic route mapping orthography to the mental lexicon, including stored word sounds. Assuming that TTS resulted from the reverse operation of the reading system, how did those two routes contribute to TTS? MK’s introspection was that TTS was triggered by both real words, which appeared in their correct orthography, and by pseudowords, which appeared in a plausible spelling, suggesting that both reading routes were feeding the TTS phenomenon. The left SMG appeared as an obvious candidate for phonological processing, considering its involvement in a broad variety of phonological tasks, particularly whenever phonology has to be interfaced with orthography, audiovisual language, or sensorimotor processing (Binder, 2017; DeWitt and Rauschecker, 2012; Hickok and Poeppel, 2016; Martin et al., 2016; Pillay et al., 2014; Savill et al., 2019). The aim of Experiment 3 was to identify regions subtending lexical access from speech in MK, allowing us to then study their contribution to TTS. To this end, MK was scanned while he was presented with spoken words and with matched pseudowords. A group of controls were also scanned (see Methods).

As usual, both words and pseudowords triggered TTS. In MK, words and pseudowords activated essentially identical left-predominant fronto-temporo-parietal areas relative to rest. Those areas encompassed the entire TTS network, as identified in Experiment 1 (Supplementary Figure S4). Unless stated otherwise, all further analyses of this experiment were restricted to the volume which was activated in MK by averaged words and pseudowords > rest (voxelwise p<0.01). We then compared MK minus controls for the contrasts words > rest and pseudowords > rest. Both comparisons replicated the results of Experiment 1, revealing overactivation in MK’s TTS network, including the IFG, SMG and SMA (Supplementary Figure S5). The VWFA also was overactivated in MK for words > rest when correcting within the same a priori ROI as in Experiment 1.

We then compared activations by words and pseudowords, using univariate contrasts and multivariate pattern analysis (MVPA). The contrasts of words > pseudowords and pseudowords > words showed no significant effect of lexicality in MK. Controls showed stronger activation for pseudowords > words in bilateral superior temporal sulci and in bilateral inferior frontal gyri. MK did not differ from controls in either of those contrasts.

Moreover, we compared activation by words and pseudowords in MK at the peak voxels of the 5 regions of the TTS network as identified in Experiment 1. MK had stronger activations for pseudowords>words in the precuneus (p<0.001), and no difference in the other regions (all Ps>0.07).

We then used an MVPA searchlight to decode activation by words vs. pseudowords. In MK, the bilateral left-predominant posterior middle temporal gyrus (MTG) and the VWFA (MNI −48 −46 −19) distinguished words from pseudowords better than chance (Figure 4). This pattern did not differ between MK and controls.

**Figure 4.**
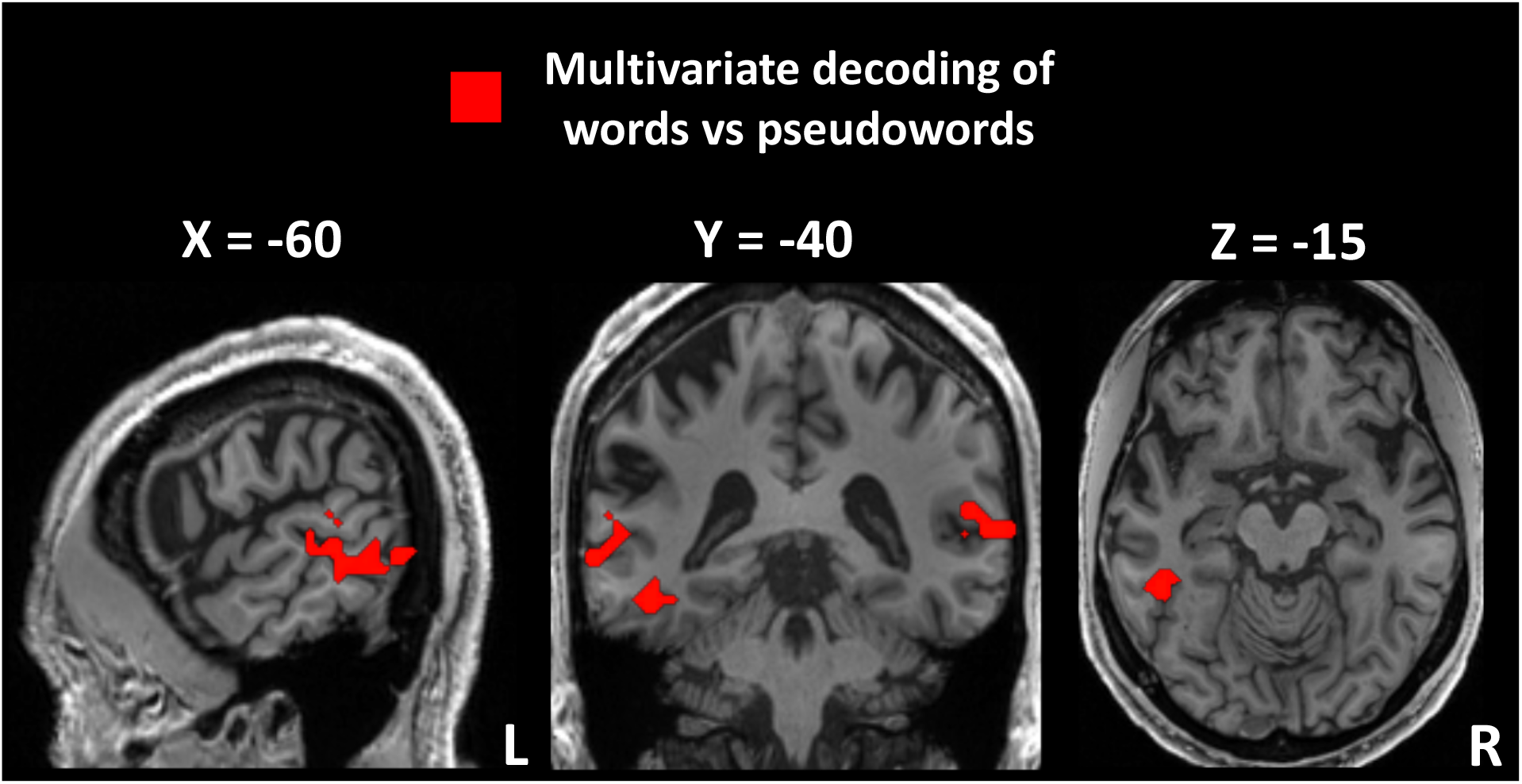
Experiment 3: Tickertape synesthesia (TTS) for words and pseudowords. MVPA discrimination between words and pseudowords in MK, showing effective decoding of lexical status in the left-predominant MTG and in the VWFA. MVPA, multivariate-pattern analysis; MTG, middle temporal gyrus; VWFA, visual word form area.

In summary, while univariate analyses showed no significant effect of lexicality in MK, MVPA showed effective decoding in the VWFA, and in posterior MTG areas. Neuropsychology and imaging studies have long involved the left posterior MTG in lexical processing (see for instance Binder et al., 2009; Boatman, 2000; Davey et al., 2016; Hart and Gordon, 1990; Pillay et al., 2014), which makes it a sensible implementation of the lexical component of TTS. Having identified a set of candidate regions involved in all aspects of TTS, we are now in a position to investigate their functional interactions.

### Dynamic causal modelling of ticker-tape synesthesia

The set of regions identified in Experiments 1-3 as associated to TTS generation belong to the usual cerebral reading network. This overlap was specifically demonstrated in Experiment 2 (Figure 3A). In addition to a higher univariate activation level in those regions, we predicted that TTS should be associated with an atypical flow of information among them, featuring an increase in top-down influences. We assessed this prediction using DCM, a method whose aim is to estimate the functional coupling among brain regions, and the changes in this coupling that are associated with different experimental conditions. Alternative models of how observed time series were generated in the brain are compared using a Bayesian approach (see Methods; Friston et al., 2003).

We modeled the causal links among the core regions involved in TTS generation, based on the hypothesis that speech perception triggers interacting phonological and lexical processes, eventually generating a vivid orthographic image. This is equivalent to assuming, as proposed before, that TTS results from the top-down operation of a two-route reading system. We re-analyzed MK’s data from Experiments 1 and 3 using dynamical causal modeling (DCM), in order to identify the best model among alternative candidates. We translated our specific hypotheses into a set of constraints on the space of candidate models.

Among the regions identified in Experiments 1-3, we selected as regions of interest the mid-STG, the SMG, the MTG, and the VWFA (see Methods). To define the space of possible models, we assumed (1) that the mid-STG is the input region for all auditory stimuli, and that it is unidirectionally connected to the SMG, the MTG, or both (3 possible configurations); (2) that the VWFA is the output region subtending orthographic representations, and that it receives unidirectional input from the SMG, the MTG, or both (3 possible configurations); (3) that the SMG and MTG are mutually connected. Combining the 3 input and the 3 output options resulted in 9 families of models (Supplementary Figure S6) (Penny et al., 2010).

We then allowed some of the connections defining these families to be modulated by experimental factors. In Experiment 1, the connections from the mid-STG to the SMG and MTG could be modulated by the normal vs reversed speech factor. This is because this difference should only be relevant early in the flow of information, i.e. low-level acoustic stimuli should not propagate to high-level speech and language representations. In Experiment 3, all connections could be modulated by the real words vs pseudowords factor (Supplementary Figure S6).

In summary, this resulted in 9 model families, including a total of 24 models for Experiment 1, and 256 models for Experiment 3.

For Experiment 1, the comparison of the 9 families selected a structure including all possible connections (posterior probability 0.86, Figure 5A). The second-best family did not include the connection from the SMG to the VWFA (posterior probability 0.11). For Experiment 3 also, the model featuring all possible connections was the most accurate (posterior probability 0.99, Figure 5B).

**Figure 5.**
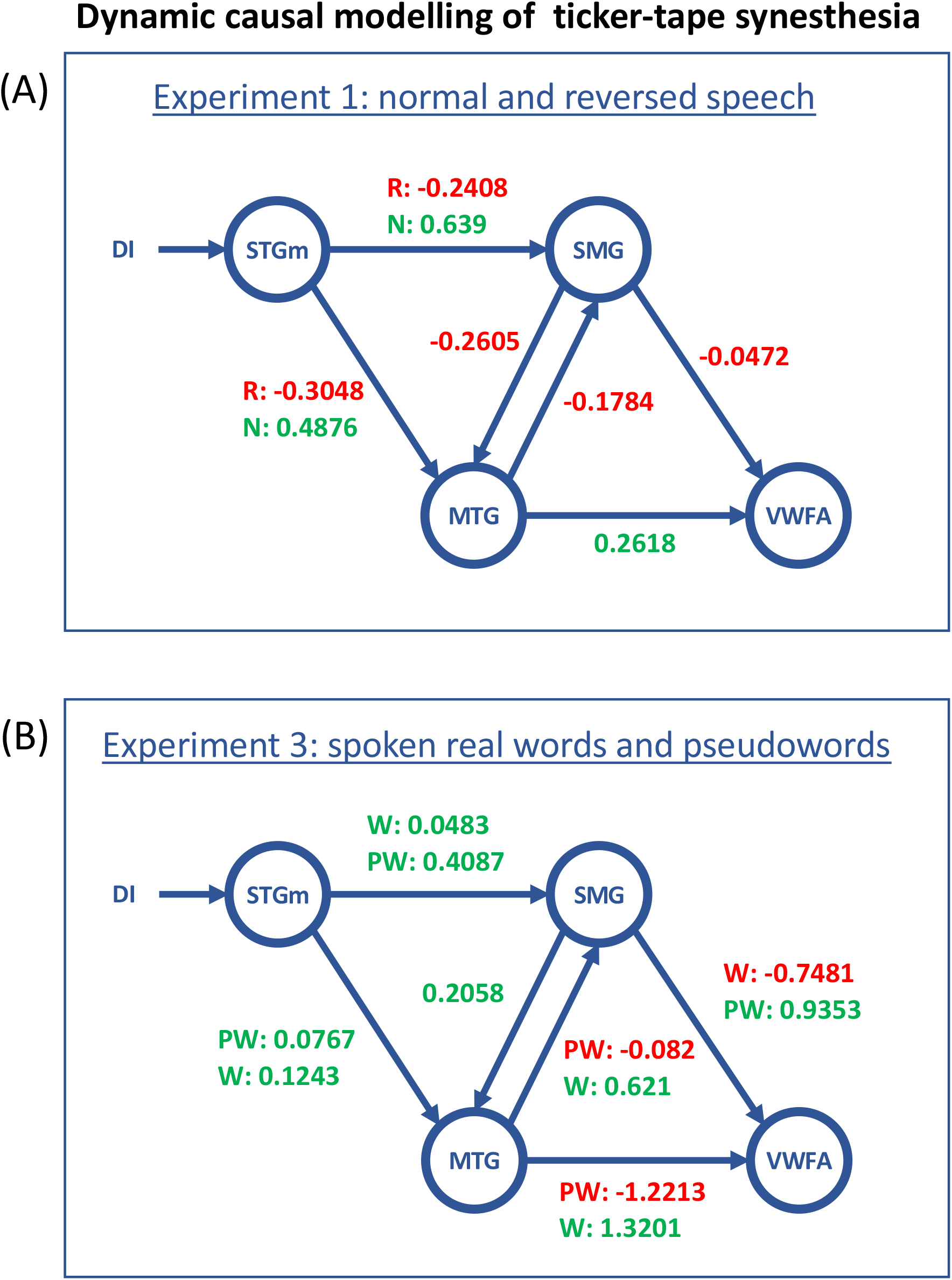
Dynamic causal modelling of ticker-tape synesthesia (TTS). Most likely models for MK’s TTS-associated activations. The driving input (DI) corresponds to auditory stimulations entering the network through the STG. Blue arrows represent the intrinsic connections between the nodes. Values represent the strength of connections, which may be positive/excitatory (green) or negative/inhibitory (red). Two values are associated to connections significantly modulated by conditions, one value corresponding to each of the two experimental conditions. (A) In Experiment 1, normal speech (N) was driving the whole network down to the VWFA, while reversed speech (R) did not. (B) In Experiment 3, connections implementing the phonological route (STG-SMG and SMG-VWFA) were excitatory for pseudowords (PW) and inhibitory for real words (W). Conversely connections involved in the lexical route (MTG-VWFA and MTG-SMG) switched from inhibitory with pseudowords to excitatory with real words. STG, superior temporal gyrus; SMG, supra-marginal gyrus; VWFA, visual word form area; MTG, middle temporal gyrus.

We then compared the models within the two winning families, in order to identify for each experiment the model with the most accurate pattern of modulation. In Experiment 1, the best model among 4 candidates had a posterior probability of 1. The connections from the STG to both the SMG and the MTG were significantly modulated, switching from strongly excitatory for normal speech to mildly inhibitory for reversed speech (Figure 5A). Regarding nonmodulated connections, on average across normal and reversed speech, the links from the MTG to the VWFA were excitatory, while reciprocal connections between the SMG and MTG and connection from SMG to the VWFA were slightly inhibitory.

In Experiment 3, the best model among 64 candidates had a posterior probability of 0.31 (Figure 5B). Connections from the STG to both the SMG and MTG were excitatory for words and pseudowords. The connections implementing the phonological route, i.e. from the STG to the SMG and from the SMG to the VWFA, were both significantly modulated, both with an advantage for pseudowords over words. Particularly, the connection from to SMG to the VWFA switched from strongly excitatory for pseudowords to strongly inhibitory for words. Connections implementing the lexical route, i.e. from the STG to the MTG and from the MTG to both the VWFA and the SMG, were also modulated, all with an advantage for words over pseudowords. Particularly, the connection from the MTG to the VWFA switched from strongly excitatory for real words to strongly inhibitory for pseudowords. On average across words and pseudowords, the unmodulated connection (SMG to MTG) was excitatory (Figure 5B). The second-best model had a posterior probability of 0.28. The modulations were the same as in the best model, minus the one from the STG to MTG. Modulations for words and pseudowords on other connections were similar. The third best model had a posterior probability of 0.20. It was also similar to the best one, with an additional modulation of the SMG-to-MTG connection. Connections of the phonological route, i.e. from STG to SMG and from SMG to MTG, still showed an advantage for pseudowords (excitatory) over words (inhibitory). Thus the second- and third-best models preserved the main features of the dual-route pattern of connection, supporting the stability of our findings.

## Discussion

### Summary of findings

The main outcome of Experiment 1, in which we presented normal and reversed speech, was that a set of areas were more active in MK than in controls during normal than reversed speech, that is whenever MK was subject to synesthesia. Supporting our first prediction, this “TTS network” included both frontoparietal areas involved in speech processing and the VWFA subtending orthography (Figure 1). In this experiment, one group of controls was not matched to MK in age, while the other group was scanned on a different device. However, this methodological imperfection cannot account for the differences between MK and controls.

First, those differences were almost perfectly replicated when comparing MK separately to both groups, clearly supporting the robustness of the effect (Figure S1). Second, the differences we found between the two groups are remote from all components of the “TTS network”, and are likely a correlate of aging (Figure S2). Thus, increasing age is associated with stronger prefrontal activation in a variety of tasks, including language tasks (for a review see Davis et al., 2014), and with weaker activation of auditory regions during speech perception (for a review see Cardin, 2016).

In Experiment 2, we delineated MK’s reading areas by comparing activation by written words minus other categories of visual items. The set of reading-specific areas turned out to be essentially identical to the TTS network from Experiment 1, thus backing our second prediction that TTS and reading should rely on the same pathways (Figure 3). In Experiment 3, we refined our approach by presenting MK with spoken words and pseudowords. Both triggered TTS and activated the TTS network, although possibly with different underlying dynamics. This experiment allowed us to identify an MTG area involved in lexical reading, and presumably in the lexical component of TTS (Figure 4). Finally, we developed a dynamic two-route model of information flow during TTS. In MK, we found that speech was driving the TTS network, while reversed speech, which does not trigger synesthesia, had an inhibitory effect (Figure 5A). Moreover, functional connections implementing the phonological route were inhibitory when synesthesia was triggered by real words, while they were excitatory when synesthesia was triggered by pseudowords. The converse pattern prevailed for connections implementing the lexical route (Figure 5B). This result fits our fourth prediction that, analogous to reading processes, top-down information should follow distinct paths depending on whether synesthesia was triggered by words or pseudowords.

We will first discuss the two major components of TTS generation: The VWFA and its role in orthographic imagery, and the perisylvian areas and their role in translating speech to orthography.

### The VWFA and orthographic mental imagery

We hypothesized that the orthographic content of TTS is supported by the VWFA, because there is ample evidence from brain damage and imaging that the VWFA computes orthographic representations from visual input. Importantly, in the absence of written stimuli, the VWFA may also be activated from top-down by speech (Cohen et al., 2021; Ludersdorfer et al., 2016; Yoncheva et al., 2010). We reasoned that such top-down effects may contribute to orthographic imagery, and that TTS may be seen as an exceptionally vivid and automatic form of such imagery, an ability possessed to some degree by all literate individuals (Bourlon et al., 2009; Holm et al., 2015). Supporting the role of the VWFA in orthographic imagery, at least some of the alexic patients with VWFA lesions lose this type of imagery (Bartolomeo et al., 2002; Rosazza et al., 2007; Sirigu and Duhamel, 2001; Volpato et al., 2012). This fits with the present findings that the VWFA was involved in TTS: the VOT region which was over-activated during TTS, and the VWFA as defined by its preference for written words, were precisely identical. In MK, TTS largely escaped voluntary control, while mental imagery is usually optional and effortful (Mazard et al., 2002). However the degree of automaticity of orthographic imagery is variable across individuals, rather than distributed in an all-or-none manner between TTS and non-TTS individuals. Thus, Holm et al. (2015) asked participants from a general population to specify the extent of their control of word visualization. A small number of subjects qualified for TTS in the strictest sense, similar to MK, synesthesia being obligatory, and induced by other persons’ speech, one’s own speech, and when thinking verbally. Interestingly, a larger number of participants reported that word visualization was under some voluntary control, relevant to the issue of continuity between synesthesia and normal perception, a controversial issue in the field of synesthesia studies (Deroy and Spence, 2016; Martino and Marks, 2001).

Considering the vividness and automaticity of mental imagery in MK, we may draw a parallel between TTS and the visual hallucinations that occur in schizophrenia, in Parkinson’s disease and other neurological conditions, and in the Charles Bonnet syndrome. Although the mechanisms of such hallucinations are still elusive, they are often associated with overactivation of visual cortices, which show abnormal intrinsic and distant connectivity, supporting predominant top-down over bottom-up influences on visual experience (Amad et al., 2014; Dujardin et al., 2020; ffytche et al., 1998; Hahamy et al., 2021; Waters et al., 2014).

### From speech to orthography

#### The superior temporal gyrus

In the dynamical model of TTS, we treated the mid-STG cortex as a common auditory input region, because it was equally activated by both speech and reversed speech. Indeed the center of the STG region of interest (MNI −48 −16 5) belonged to the early auditory cortex, as defined architectonically, with a probability of 85% (areas TE 1.0 1.1 1.2; Morosan et al., 2001). Further supporting the role of the mid-STG as the initial stage of TTS generation, DCM showed that it was significantly driving the SMG and MTG whenever MK was subject to synesthesia.

In MK, both externally perceived and inner-speech triggered synesthesia. This may result from the remarkable overlap of STG activation by external and by inner-speech (Alderson-Day et al., 2016; Alderson-Day and Fernyhough, 2015; Shergill et al., 2002), and also during auditory-verbal hallucinations in schizophrenia patients (Jardri et al., 2011).

#### The supramarginal gyrus

Beyond the auditory entry point, the STG is a complex mosaic of modules computing various representations of speech and sounds (Binder, 2015; DeWitt and Rauschecker, 2012; Scott, 2019; Yi et al., 2019). In continuity with the posterior STG, the SMG supports phonological representations at the interface with other modalities. This includes audio-motor mapping for speech production (Hickok and Poeppel, 2015), phonological working memory (Buchsbaum et al., 2005; Buchsbaum and D’Esposito, 2008), lip-reading and audiovisual language (e.g. Dick et al., 2010; Hocking and Price, 2008; Wright, 2003).

It is also critically involved in non-lexical reading, such that per-operative stimulation of the SMG disrupts selectively the reading of pseudowords but not of exception words (Roux et al., 2012; Simos et al., 2000). Similarly, transcranial stimulation of the SMG interferes with sound-oriented but not with meaning-oriented reading (Pattamadilok et al., 2010; Sliwinska et al., 2015). Further supporting its role in phoneme-grapheme correspondences, the SMG is more activated by pseudowords than real words, both during reading (Taylor et al., 2013) and during spelling (DeMarco et al., 2017). Graves & Binder (2010) found that the left SMG showed a negative effect of bigram frequency, millimeters away from the current SMG peak, revealing sensitivity to the difficulty in binding phonemes and graphemes. Moreover Bouhali et al. (2019) found stronger functional connectivity of the SMG with the VWFA when reading aloud pseudowords than during lexical decision with words, that is whenever phonological demands were higher.

In MK, the role of the SMG in TTS may be to map perceived phonemes to illusory graphemes on the basis of statistical regularities, in interaction with the lexical route. One may putatively view the overactivation of the SMG in synesthesia as functionally symmetric to the hypoactivation observed repeatedly in developmental dyslexia, where phoneme-grapheme mapping is impaired (D’Mello and Gabrieli, 2018; Kronbichler et al., 2006).

#### The middle temporal gyrus

The posterior MTG is one of the most systematically activated regions during lexico-semantic tasks (Binder et al., 2009), and electrical stimulations cause transcortical sensory aphasia, that is an impairment of language comprehension with preserved phonological processing (Boatman, 2000), in agreement with the consequences of focal lesions (Hart and Gordon, 1990). This region may be seen as a “convergence hub” for binding spread-out pieces of lexical knowledge (Patterson et al., 2007), or similarly as a “semantic control” region (Davey et al., 2016, 2015; Jedidi et al., 2021).

During reading, the posterior MTG is also consistently activated when contrasting words minus pseudowords (for meta-analyses see McNorgan et al., 2015; Taylor et al., 2013), supporting its involvement in the lexical reading route (Binder et al., 2003). However, lexicality interacts with task, such that MTG activation is mostly present during tasks with strong lexical demands, typically lexical decision relative to reading aloud (McNorgan et al., 2015). This may explain why in Experiment 3, where participants only had to detect an odd-ball meaningless signal, we found no univariate effect of lexicality. In the MTG, MVPA proved more sensitive for discriminating words from pseudowords, similar to previous findings (Baeck et al., 2015; Fedorenko et al., 2012; Mattheiss et al., 2018).

This background made the MTG, as identified in MK during synesthesia, a sensible implementation of lexical processing within a simple model of TTS.

After discussing the putative roles of the VWFA and perisylvian areas in TTS, we now turn to the dynamic model of their interactions which we explored using DCM.

### A simple dynamic model of TTS

#### Constraints on the model

When designing the DCM model of synesthesia, we selected a set of 4 areas of interest, admittedly a simplistic network, leaving aside the IFG, SMA and precuneus, although they were also over-activated in MK. This allowed us to restrict the space of possible models on the basis of strong a priori motivations, a methodological requirement of DCM (Stephan et al., 2010). The STG, SMG, MTG, and VWFA are essential components of auditory, phonological, lexical, and orthographic processing, respectively, and were all defined on the basis of their functional features during TTS. The 4 selected regions thus provided a sensible sampling of the lexical and phonological routes.

We reasoned that the IFG and SMA probably played no critical role in the generation of TTS. While the IFG is often activated by reading words and pseudowords (e.g. Bouhali et al., 2019; Heim et al., 2005), it is thought to control spoken phonological output, and should not contribute to the purely audio-visual phenomenon of TTS (Taylor et al., 2013). The SMA is also commonly activated during reading, and its primary language-related role is thought to be the control of speech output (Alario et al., 2006; Hertrich et al., 2016). Finally, the precuneus is a top-level hub in the hierarchy of brain areas, it intervenes in a variety of cognitive control processes, and its putative role in TTS is difficult to pinpoint from the current data (Cavanna and Trimble, 2006; van den Heuvel and Sporns, 2013). Still in a recent meta-analysis, Spagna et al. (2021) proposed that activation of both the SMA and precuneus are associated with mental imagery, and their putative role in TTS may deserve further study.

In defining the space of possible models, we posited that the SMG and MTG, as implementing the phonological and lexical routes, should always have the possibility of influencing each other. There is a host of behavioral evidence of close interactions between the two routes. Thus, illustrating the automatic impact of phonology on lexical reading, during visual lexical decision, words are automatically parsed into phonologically defined syllables (Conrad et al., 2007). In the same task, response to word targets is facilitated by pseudoword homophone primes (mayd MADE), as compared to control primes (mard MADE), revealing the computation of phonological codes and their impact on word reading (Grainger and Holcomb, 2008). Conversely, showing the influence of lexical knowledge on phonological reading, pseudowords are pronounced faster when they are orthographically similar to a large number of real words (Peereman and Content, 1995). A point of interest in the context of TTS as “inverted reading”, not only the reading but also the spelling of pseudowords is also influenced by the existence and frequency of real word neighbors (Tainturier, 2013).

At the brain level, close interactions between the SMG and posterior MTG are supported by their strong anatomical connections (Caspers, 2011; Jung et al., 2017; Xu et al., 2016) and their functional connectivity at rest (Turken and Dronkers, 2011; Xu et al., 2016).

#### Architecture and modulations of the TTS network

In Experiment 1, normal speech had a strong excitatory driving influence on the circuit, while reversed speech had a minor inhibitory effect. Moreover, in the most likely model family, words propagated from the STG to both the MTG and SMG, while only the latter further relayed information to the VWFA. In other terms, until they are assigned an orthographic form by the VWFA, perceived words mostly follow the lexical route through the MTG. The specific contribution of the MTG here may be to allow access to stored orthographic knowledge, possibly by performing semantic disambiguation of auditory words (Davey et al., 2016, 2015; Jedidi et al., 2021; Patterson et al., 2007). Finally, the phonological route (SMG) only had a small inhibitory effect on the lexical route (MTG), suggesting that it did not play an important role in generating TTS for normal speech.

This interpretation is aligned with the DCM results of Experiment 3 which, in contrast to Experiment 1, included both normal speech (real words) and pseudowords. The most likely network included the same connections as in Experiment 1. When first neglecting inhibitory effects, Experiment 3 essentially recapitulates Experiment 1, in that real words reach the VWFA through the lexical route, i.e. through the MTG. In contrast, pseudowords drive the VWFA through the phonological route, i.e. through the SMG. The pattern of inhibitory effects further suggests that the lexical route (MTG to VWFA) acts to lessen VWFA responses to pseudowords, whereas the phonological route (SMG to VWA) dampens VWFA responses to real words. This indicates that the two routes are somehow competing for access to the VWFA, eventually facilitating accurate orthographic coding of both words and pseudowords. Overall, these results support our fourth hypothesis that top-down influences should follow distinct lexical vs phonological paths depending on whether synesthesia is triggered by words or pseudowords.

In Experiment 3, we found only a weak excitatory effect of real words from the STG to the SMG, while in Experiment 1 the excitatory effect of normal speech was strong. As real words consist in normal speech, we may have expected a strong positive effect of words in Experiment 3 also. Leaving aside DCM reproducibility issues, this discrepancy may result from differences in task requirements: In Experiment 1, participants payed attention to a meaningful narration while in Experiment 3, they were encouraged to process words at a low level. Words lists carried no sentential meaning, and the target “tatatata” could be best detected by paying attention to phonology. This may explain why words had a stronger influence in Experiment 1 than 3. Direct comparisons of words, pseudowords and low-level sounds with a constant task could clarify this issue.

Schematically, we construed TTS as an idiosyncratic operation of the reading system. We will now briefly discuss the alternative view of TTS as inner spelling.

### TTS as inner spelling

We proposed that TTS resulted from intense top-down activation of the reading network, initially because MK reported a subjective experience of inner reading, with no active spelling-related feeling. Alternatively however, TTS could have been construed as an extreme form of inner spelling. In some sense MK was automatically “writing out in his head” the speech he was perceiving. Actually those two points of view are largely equivalent, as writing and spelling share core cognitive components and neural substrates (for reviews see Purcell et al., 2017; Rapp and Dufor, 2011). There is thus a shared set of perisylvian regions whose lesion yields both reading and spelling deficits (Philipose et al., 2007).

Focusing on regions which are presumably involved in MK’s synesthesia, the left IFG and the VWFA are activated by both reading and spelling, show cross-task adaptation suggestive of shared orthographic representations (Ellenblum et al., 2019; Purcell et al., 2017, 2011), and are both sensitive to word frequency during spelling and reading (Graves et al., 2010; Rapp and Dufor, 2011). The left SMG, posterior MTG, precuneus and SMA are also activated during both reading and spelling (DeMarco et al., 2017; Rapp and Dufor, 2011). Moreover, the SMG plays a predominantly phonological role during spelling, as it does during reading and in TTS (DeMarco et al., 2017). Thus, TTS may reflect atypical functioning, during speech perception, of the single network which links sounds and letters during both reading and writing.

### Ticker tape synesthesia and brain connectivity

Beyond TTS, atypical anatomical or functional connectivity is at the core of most theories of synesthesias (for a recent review see Ward and Simner, 2020). Within this broad framework, theories differ along multiple dimensions: is there an increase in anatomical or functional connections, are those disinhibited or structurally stronger, do they link the “inducer” and “concurrent” regions directly or through higher-level cortices, do the same principles apply to all forms of synesthesias, etc? For instance, in grapheme-color synesthesia, the most studied type of synesthesia, increased fractional anisotropy was detected in the fusiform and other regions (Rouw and Scholte, 2007). Furthermore, differences in functional connectivity were found between synesthetes and controls (Sinke et al., 2012), and between subtypes of grapheme-color synesthetes (van Leeuwen et al., 2011). However the reproducibility of those findings across studies appears questionable (Hupé and Dojat, 2015). Our DCM analysis clarifies the pattern of functional connectivity involved in TTS, but is not in a position to adjudicate between general theories of synesthesia.

Of particular interest is the idea that synesthesia “magnifies connections present in early life that are pruned and/or inhibited during development and that persist in muted form in all adults” (Spector and Maurer, 2009). In this hypothesis, TTS would reflect the persistence of a transient state of stronger connection which would normally vanish after literacy acquisition. Indeed, in a longitudinal study of reading acquisition, Dehaene-Lambertz et al. (2018) observed an increase in word-specific activation in parietal (MNI −44 −42 46) and frontal (MNI −48 6 24) areas during the first year of alphabetization, only millimeters away from MK’s SMG and IFG overactivation peaks. These two regions later show a decrease in activation, and are thought to play a transient role in reading acquisition, specifically in binding phonemes and graphemes. One may thus speculate that long-term survival of a normally transient role of the SMG and IFG, contributes to the genesis of TTS.

Furthermore, it is also possible that TTS involves the persistence of processes required for spelling rather than reading acquisition. Indeed effective mental imagery in children does not predict better reading acquisition, while it predicts spelling acquisition (Guarnera et al., 2019). During spelling, it is necessary to generate and maintain an orthographic representation of the current word while serially writing or typing the component letters (Buchwald and Rapp, 2004).

This orthographic working memory, or “graphemic buffer”, may involve strings of abstract letters, but also visual images of the current string, providing a target pattern and allowing for feedback control during the writing process. Visual control plays a limited role in expert writers (Danna and Velay, 2015), but it is critical during handwriting learning (Meulenbroek and Van Galen, 1988). Thus TTS might be construed as the persistence of an hyperactive graphemic buffer and its associated visual images. Indeed the maximum overlap of lesions yielding graphemic buffer impairments is millimeters away from MK’s SMG over-activation (Rapp et al., 2016), while the VWFA would support mental images per se as discussed before.

As a final topic for speculation related to the developmental origins of TTS, one may mention the intriguing case of a man in whom TTS was associated with two other developmental conditions, chronic psychosis and an arachnoid cyst (Bastiampillai et al., 2014).

### Conclusion

Among the many kinds of synesthesia, as many as 88% involve language either as the “inducer” as in grapheme-color synesthesia, or as the “concurrent” as in perfect pitch (Simner, 2007; Simner et al., 2006). TTS stands out as the single form of synesthesia in which both the inducer and the concurrent are language representations. Specifically, TTS involves the same orthographic and phonological representations which support reading and spelling. One may thus speculate that TTS results from an atypical developmental trajectory with enduring hyperconnectivity between speech and vision, a situation in some sense symmetrical to developmental dyslexia, in which defective connectivity was repeatedly observed (e.g. Cui et al., 2016; Schurz et al., 2015; van der Mark et al., 2011).

At this stage, the present study being restricted to a single case, and in the absence of any previous data on the cerebral mechanisms of TTS, all general inferences should be cautious. Francis Galton (1883) noted that “the experiences differ in detail as to size and kind of type, colour of paper, and so forth, but are always the same in the same person”, pointing to the issue of individual variability. Beyond the perceptual features of TTS mentioned by Galton, synesthetes also differ in the level of control that they exert on TTS, and in the stimuli (external speech, inner speech, etc) that trigger the phenomenon (Holm et al., 2015). We predict that all cases of TTS should conform to the general pattern of “upended reading”, but expect that the exact pattern of visual activation, and the involvement of attention and control systems should vary substantially.

Further study of TTS may therefore shed light on the mechanisms of reading acquisition, their variability and their disorders. Among others, future studies should address resting-state and anatomical connectivity in TTS individuals, potential behavioral differences in oral and written language processing between TTS individuals and controls, associations with other synesthesias, variants and dissociations within TTS, and TTS-like phenomena in typical and dyslexic children learning to read.

## DATA SHARING PLANS

Data are available at https://zenodo.org/record/6900980.

