## Supplementary Figure S1, S2, S3, S4, S5, S6, Table S1, S2 for "Seeing speech: The cerebral substrate of tickertape synesthesia"

### Supplementary information


- Supplementary Figure S1
- Supplementary Figure S2
- Supplementary Figure S3
- Supplementary Figure S4
- Supplementary Table S1
- Supplementary Table S2

Supplementary Figure S1

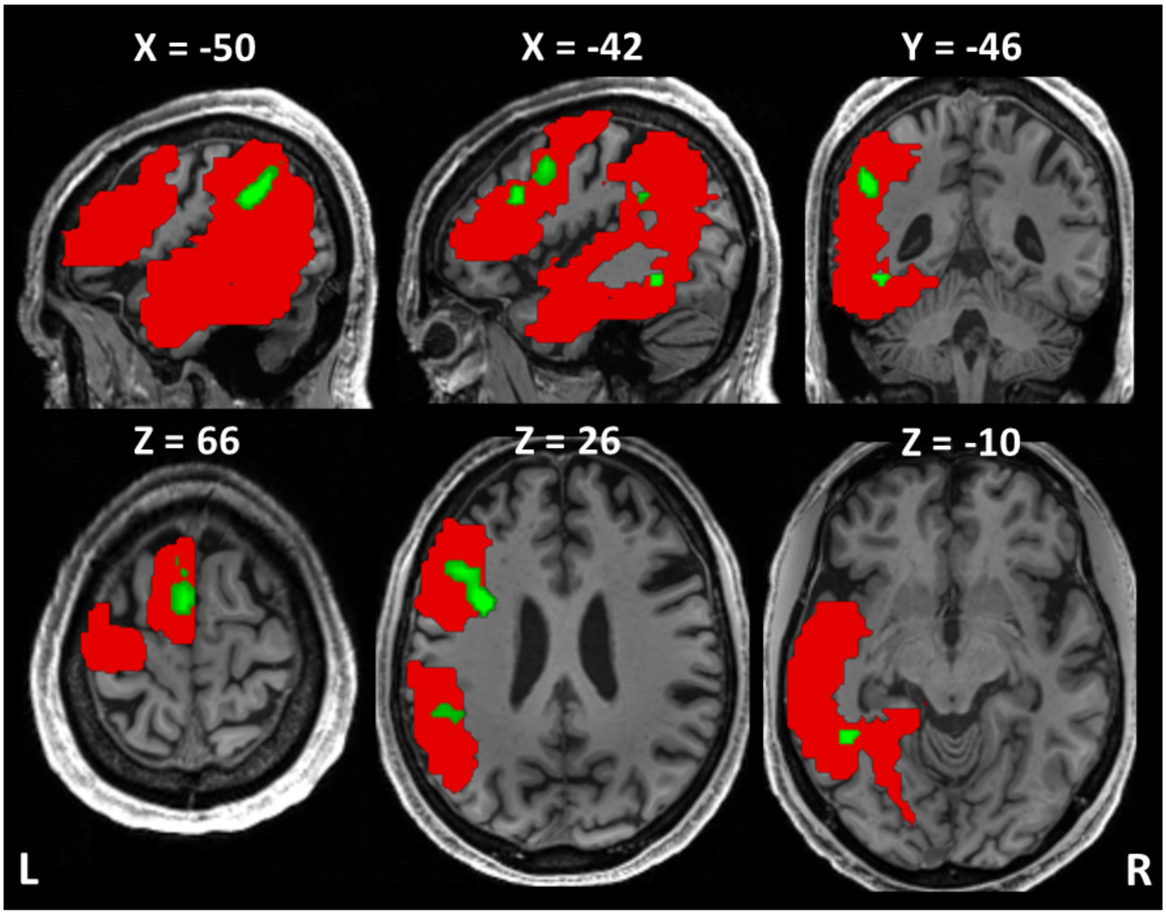


Anatomical mask of language network

Normal > reversed speech in MK > controls

For some analyses of **<u>Experiment 1</u>**, we defined a broad region of interest covering left-hemispheric language areas and VOT cortex (red), by merging the opercular and triangular parts of the inferior frontal gyrus, the supplementary motor area, the precentral, fusiform, inferior parietal, supramarginal, angular, superior and middle temporal, and inferior temporal gyri, as defined in the AAL 3 atlas (Rolls et al., 2020). The "TTS network" (green) was included in this broad region of interest.

Supplementary Figure S2

(A)

(B)

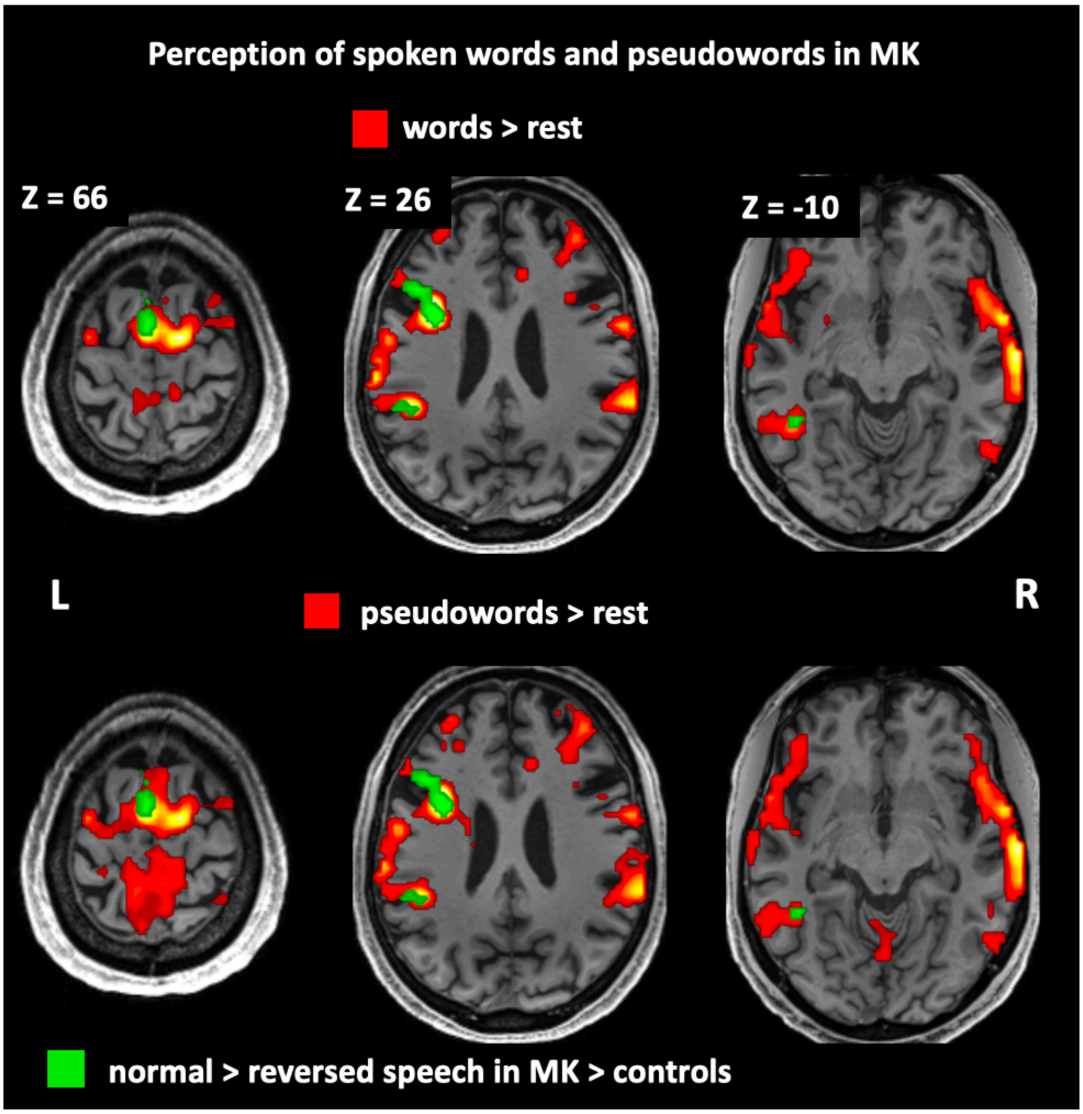


In **Experiment 3**, words and pseudowords triggered tickertape synesthesia (TTS) in MK. Activated regions for spoken words (A) and spoken pseudowords (B) relative to rest (hot colors) are essentially identical irrespective of lexicality, and encompass areas over-activated in MK during TTS in Experiment 1 (green). (C)

Supplementary Figure S3

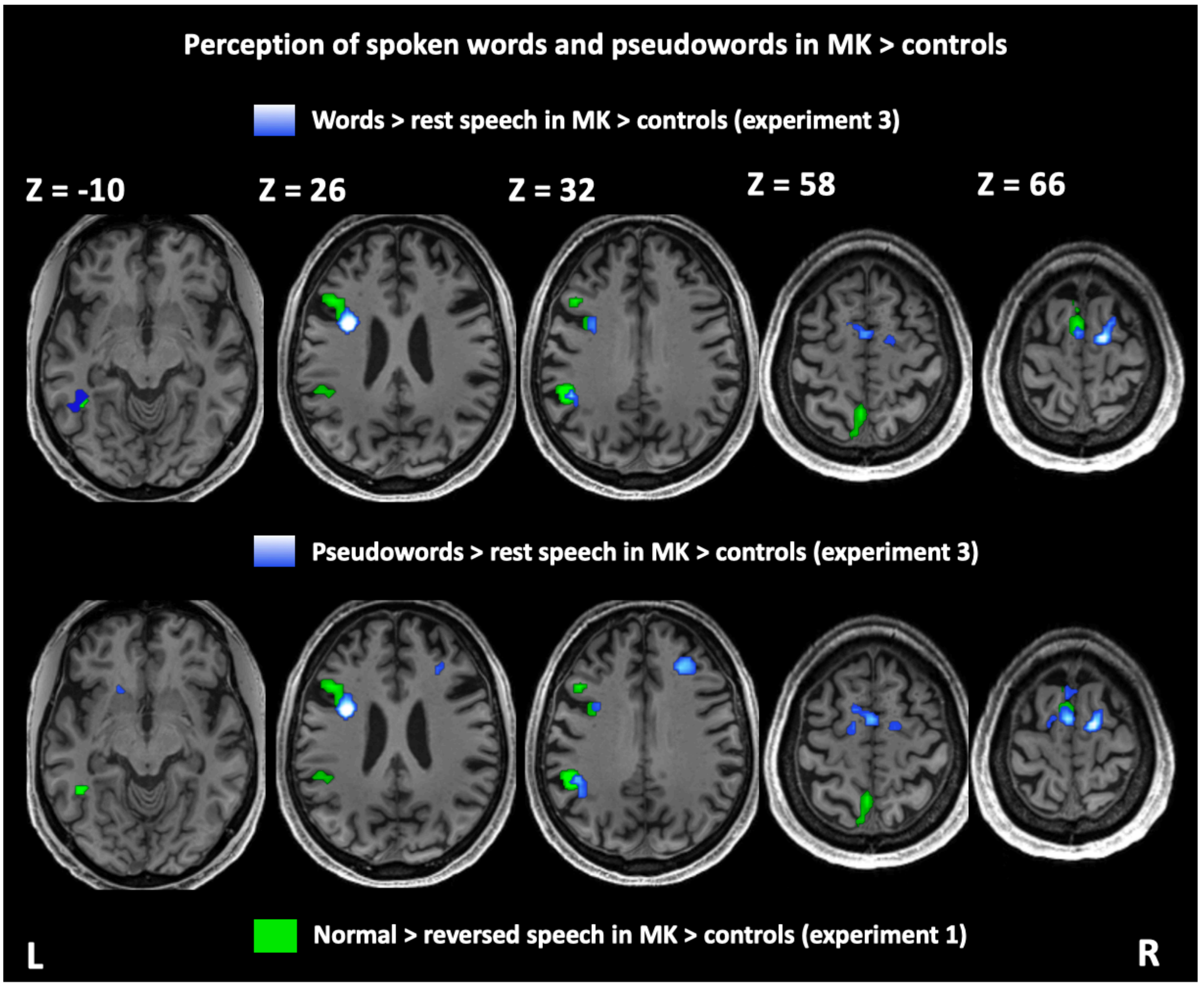


In **Experiment 3**, both spoken words and pseudowords triggered TTS in MK. The comparison of MK minus controls for the contrasts words > rest (top row) and pseudowords > rest (bottom row) revealed overactivation in MK's TTS network (blue), in regions overlapping with those observed in Experiments 1 (green) and Experiment 2 (see Figure 3).

Supplementary Figure S4

#### Model families and modulations for Dynamic Causal Modelling

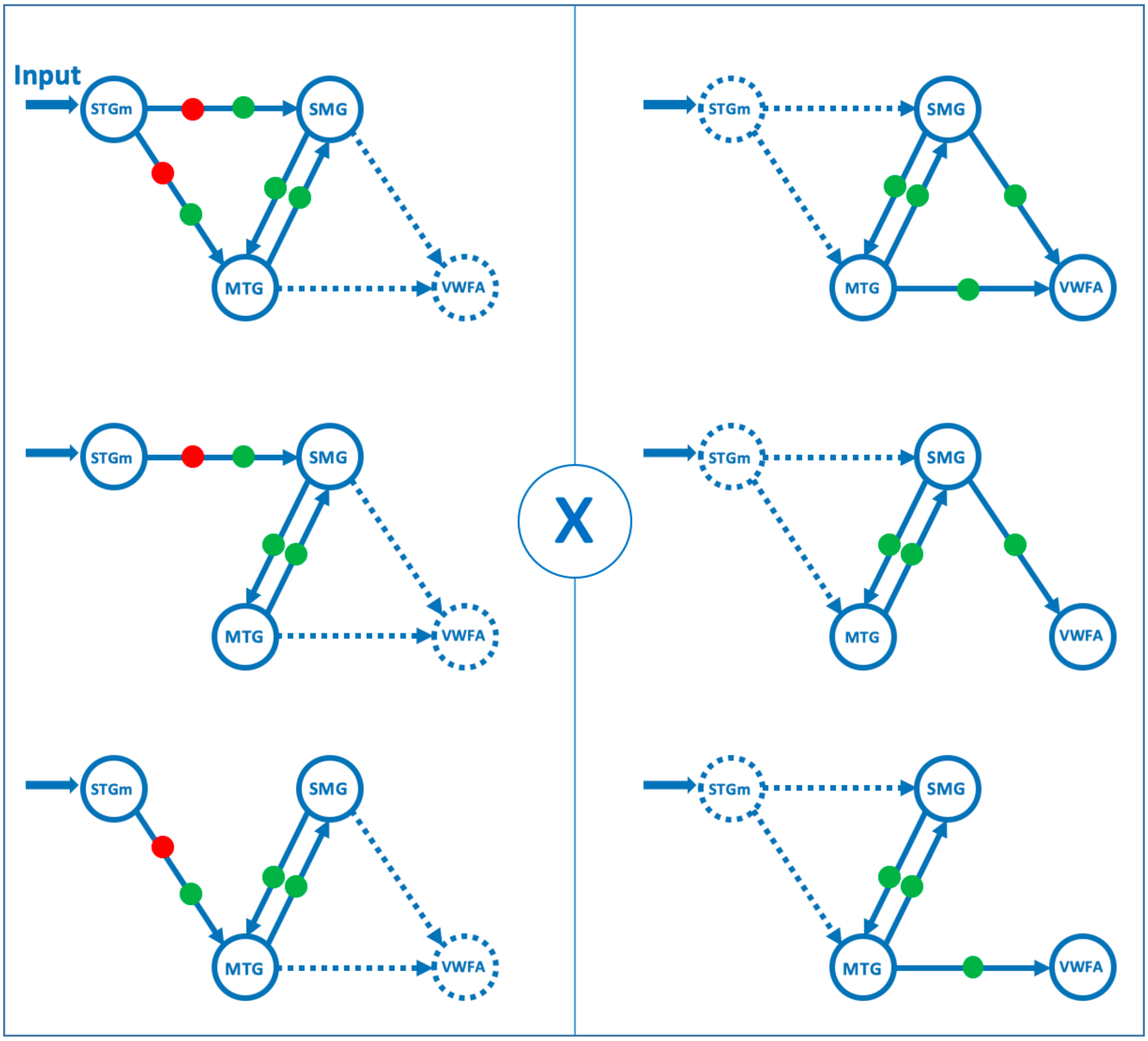


**Model families and modulations for Dynamic Causal Modelling.** All auditory stimulations drive the model through the STG (thick arrow), the SMG implements the phonological route, the MTG the lexical route and the VWFA the orthographic output. The STG drives unidirectionally the SMG, the MTG or both (3 options depicted by solid arrows in the left panel). The SMG and MTG are always reciprocally connected, implementing the interaction of the phonological and lexical routes. The VWFA receives unidirectional input from the SMG, the STG, or both (3 options depicted by solid arrows in the right panel). Combining the 3x3 options results in 9 possible families of structural models. Red dots show allowed modulations by the contrast of normal vs reversed speech (Experiment 1). Green dots show allowed modulations by the contrast of real vs pseudowords (Experiment 3).

**Supplementary Table S1.**

**Experiment 1: Regions activated in controls when listening to reversed speech, and to normal > reversed speech.**

| Reversed speech > rest | | | | Normal > reversed speech | | | |
|---|---|---|---|---|---|---|---|
| **Region** | **Peak coordinates (MNI)** | **Z-score** | **Cluster-size** | **Region** | **Peak coordinates (MNI)** | **Z-score** | **Cluster-size** |
| Left STG | -48 -16 5 | >8 | 1263 | Left STS | -54 5 -16 | 7.56 | 4195 |
| Right STG | 51 -16 8 | >8 | 1318 | Right STS | 48 14 -22 | 6.40 | |
| Right mesencephalic pedonculus | 18 -25 -4 | 5.21 | 42 | Left SMA | -39 11 53 | 5.32 | 96 |
| Right postero-inferior frontal gyrus | 42 11 20 | 5.16 | 211 | Right inferior frontal | 54 26 5 | 4.76 | 50 |
| Right SMA | 51 2 50 | 4.71 | 51 | Bilateral pre-cuneus | -6 52 38 | 4.73 | 177 |
| Left postero-inferior frontal gyrus | -33 14 29 | 4.04 | 48 | Left medial SMA | -6 5 65 | 4.52 | 50 |
| | | | | Bilateral orbitofrontal cortices | 0 59 -16 | 4.35 | 38 |

MNI, Montreal Neurological Institute; STG, Superior temporal gyrus; SMA, Supplementary motor area; STS, Superior temporal sulcus.

**Supplementary Table S2.**

**Experiment 2: Regions showing preferential activations for categories of visual stimuli.**

| Contrast | Region | Peak coordinates (MNI) | $Z_{max}$ |
|---|---|---|---|
| **Words > Houses and Faces** | Left STG | -63 -22 5 | > 8 |
| | Left fusiform gyrus | -45 -52 -10 | > 8 |
| | Right STG | 69 -31 11 | 7.71 |
| | Left inferior frontal gyrus | -33 8 23 | 6.85 |
| | Left supramarginal gyrus | -48 -40 32 | 6.29 |
| | Left precuneus | -9 -73 59 | 4.72 |
| | Right cuneus | 12 -97 14 | 4.6 |
| | Left SMA | -6 8 62 | 4.43 |
| **Faces > Houses and Words** | Right peristriate area | 51 -70 -1 | 6.38 |
| | Right fusiform gyrus | 42 -55 -13 | 5.44 |
| **Houses > Faces and Words** | Right visual association area | 30 -88 8 | > 8 |
| | Left visual association area | -33 -85 23 | > 8 |
| | Left parahippocampal gyrus | -30 -37 -16 | > 8 |
| | Right parahippocampal gyrus | 30 -49 -10 | 6.09 |
| **Tools > (Words, Houses and Faces)** | Left lateral occipital cortex | -42 -67 2 | > 8 |
| | Right lateral occipital cortex | 45 -64 -4 | > 8 |
| | Left fusiform gyrus | -30 -49 -13 | 5.08 |
| **Bodyparts > (Words, Houses and Faces)** | Right lateral occipital cortex | 45 -64 -4 | > 8 |
| | Left lateral occipital cortex | -42 -67 2 | > 8 |
| | Left fusiform gyrus | -30 -52 -13 | 5.1 |

MNI, Montreal Neurological Institute; STG, superior temporal gyrus; SMA, supplementary motor area.
